# DNA methylation rates scale with maximum lifespan across mammals

**DOI:** 10.1101/2023.05.15.540689

**Authors:** Samuel. J. C. Crofts, Eric Latorre-Crespo, Tamir Chandra

**Affiliations:** MRC Human Genetics Unit, University of Edinburgh, Edinburgh, UK; School of Biological Sciences, University of Edinburgh, Edinburgh, UK; School of Informatics, University of Edinburgh, Edinburgh, UK

**Author notes:** **Correspondence to: Eric Latorre-Crespo:**, **Tamir Chandra:**. These authors contributed equally. **<u>Author Contributions:</u>** E.L.C. and T.C conceived and supervised the study. S.J.C.C, E.L.C and T.C. wrote the manuscript. S.J.C.C., E.L.C and T.C. conducted data analysis. **<u>Competing Interest Statement:</u>** None applicable.

## Abstract

DNA methylation rates have previously been found to broadly correlate with maximum lifespan in mammals, yet no precise relationship has been observed. We developed a statistically robust framework to compare methylation rates at conserved age-related sites across mammals. We found that methylation rates negatively scale with maximum lifespan in both blood and skin. The emergence of explicit scaling suggests that methylation rates are - or are linked to - an evolutionary constraint on maximum lifespan acting across diverse mammalian lineages.

## Main

Organisms display enormous variation as the result of evolution, spanning many orders of magnitude in characteristics such as size, energy requirements, and lifespan. Despite this remarkable diversity, it has been observed that biological traits often share underlying mechanisms and constraints^1^. These fundamental connections between organisms can be reflected in scaling laws, which mathematically describe an association between two physical quantities over several orders of magnitude.

A notable example of a scaling law in the field of biology is Max Kleiber’s observation that an animal’s metabolic rate is proportional to its mass to the power of three-quarters^2^. This observation was later shown to hold across not just whole organisms, but also cells and mitochondria spanning a total of 27 orders of magnitude in mass^3^. It has been proposed that this relationship arises from the transport of materials through branching fractal-like networks and that evolution tends to minimise the energy required to supply these materials^4^. Such an explanation demonstrates the power of scaling laws to reveal fundamental processes that govern biological systems.

DNA methylation is an epigenetic modification in which a methyl group is added to a cytosine base followed by a guanine (CpG). Methylation status at a given CpG can vary between cells, meaning a methylation proportion can be calculated for each CpG across a population of cells. Methylation proportions of some CpGs change in a predictable way with age. This observation led to the development of the first “epigenetic clocks” in the early 2010s^5–7^ which used methylation proportions of selected CpGs to predict chronological age in humans. Since then, epigenetic clocks have been extended to numerous other organisms, including the development of clocks which measure age across mammalian species^8,9^.

Recently, in mammals, DNA methylation rates have been shown to generally correlate with a species’ maximum lifespan, although no scaling has been observed and the biological mechanisms behind the correlation remain unclear. Lowe et al. (2018)^10^ looked at age-related CpGs across six mammals and found a negative trend between methylation rates and maximum lifespan. Similarly, Wilkinson et al. (2021)^11^ looked at age-related CpGs in 26 species of bat and again found a negative correlation between methylation rate and longevity. More generally, methylation dynamics have recently been used to develop epigenetic predictors of life-history traits^12^, to attempt to identify specific CpGs and associated genes involved in both ageing and longevity^13^. Findings such as these have led to the prediction that a scaling relationship might exist between methylation rates and maximum lifespan^14^.

We compared the methylation rates of conserved age-related CpGs in blood and skin in a total of 42 mammalian species, representing 9 taxonomic orders and covering almost the entire range of mammalian lifespans (see Extended Data Tables 1 and 2). In contrast to previous studies, we remove the impact of potential statistical artefacts - that arise when comparing linear rates in a bounded space of methylation values across species of different lifespans - by developing a statistically robust framework and analysing the effect of CpG selection (described below). We found that methylation rates in both tissues scaled tightly with maximum lifespan. The emergence of explicit scaling suggests that epigenetic mechanisms are - or are linked to - an underlying evolutionary constraint on lifespan that is shared across species.

We aimed to compare methylation rates - defined as the slope from linear regressions of methylation ∼ age - in conserved age-related sites across mammals. An overview of our workflow is depicted in Fig. 1a. We initially curated our data for each tissue by removing outliers using density-based clustering^15^ on principal components (Step 1 in Fig. 1a, Extended Data Fig. 1). Additionally, we removed samples below the age of sexual maturity known to exhibit non-linear dynamics^7^ (Step 2 in Fig. 1a).

**Fig. 1:**
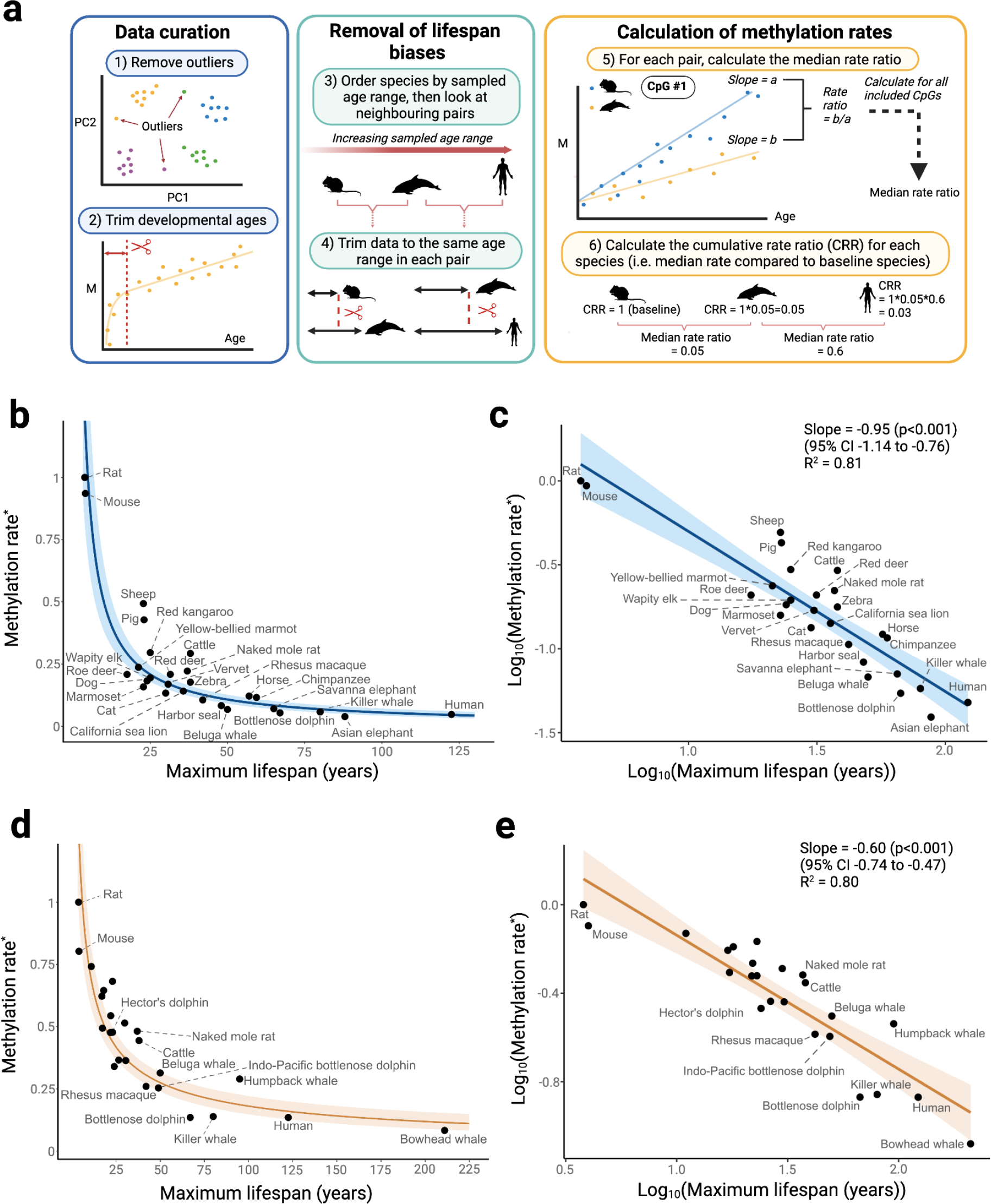
DNA methylation rates scale with maximum lifespan. **a**, Workflow overview. **b**, Methylation rate (*ratio compared to baseline species) versus maximum lifespan in blood samples. The y-coordinate of each point is the cumulative product of the median rate ratio (see Methods). Regression line plotted from the transformed log-linear association shown in c). Shaded region represents the 95% confidence interval. **c**, Same data as in b), but with axes log-transformed. Regression line from a simple linear regression of the form y∼x. **d**, and **e**, show an equivalent analysis as b) and c), but in skin samples. Unlabelled points are various species of bat (see Extended Data Table 2 for details). Created with BioRender.com.

Next, we aimed to avoid any biases arising from the calculation of rates across different lifespans. First, since methylation levels are constrained between 0 and 1, they are more likely to reach these boundaries in age-related sites and start to stabilise in longer-lived mammals. Consequently, simply fitting linear regression lines to this data will result in slower methylation rates for longer-lived animals (Extended Data Fig. 2a). Second, R^2^-based thresholds to select age-related sites may bias the selection of CpGs towards those with slower rates in longer-lived animals. This is because shorter-lived species might not be sampled long enough for small trends to become statistically apparent. Working with cohort data, these concerns arise when comparing mammalian species across different ranges of sampled ages rather than different lifespans. In a simulation based on the lifespans observed in our data, we show that not accounting for these differences in sample age ranges results in an artificial negative association with maximum lifespan (Extended Data Fig. 2b).

To develop a statistically robust framework we therefore ordered the datasets by maximum observed age and compared each mammal with its neighbours in a sequential pairwise manner (Step 3 in Fig. 1a and Methods). We started from the mammal with the shortest observed age, which we refer to as the baseline species. For each comparison, we restricted the datasets of both mammals to be as close as possible to each other (Step 4 in Fig. 1a and Methods).

For each pairwise comparison, we selected the common set of age-related CpGs passing an R^2^ threshold and sharing the same directionality. For each CpG in this set, we calculated the methylation rate for each species (Step 5 in Fig. 1a). We then calculated the methylation rate ratio of the longer-sampled species compared to the shorter-sampled species and extracted the median ratio across all selected CpGs. Then, we computed the cumulative product of median ratios to compare all species together (Step 6 of Fig. 1a, and Methods). This cumulative product can be thought of simply as the methylation rate of each species (indirectly) compared to the baseline species. In this way, we were able to compare the methylation rates of the shortest-sampled animals with the longest, while at all steps comparing rates over the same timespans.

We explored incremental R^2^ thresholds from 0 to 0.2 to define an age-related site, and show the emergence of a stable scaling law for each tissue (Extended Data Fig. 3).

To validate that our methodology removes any biases resulting in artificial scaling, we applied it to simulated data that uses the ages observed in our dataset but with rates randomly drawn for each site from all the observed rates across all mammals. The absence of any scaling law observed in this simulation emphasises the robustness of our approach in contrast to previous methods (Extended Data Fig. 4)^10,11,16^.

To explore the existence of a scaling law between methylation rate and lifespan, we plotted the methylation rate for each species (as explained above) against maximum lifespan in two tissues for which we had sufficient data - blood and skin (Fig. 1b-e). For each tissue, this revealed a relationship in which methylation rates decay to an asymptote as lifespan increases (Fig. 1b, 1d). Taking the logarithm of the x- and y-axes (see Methods for details) resulted in strong linear associations with slopes equal to -0.95 in blood (95% CI -1.14 to -0.76) and -0.60 in skin (95% CI -0.74 to -0.47) (Fig. 1c, 1e). This implies power law relationships for each tissue in which methylation rates are proportional to lifespan to the power of -0.95 (blood) and -0.60 (skin). The relationships were strong and consistent in both tissues, with relatively little variation in methylation rates unexplained by differences in maximum lifespan (R^2^=0.81 in blood, R^2^=0.80 in skin). Similar associations were seen when CpGs were stratified into hyper- and hypo-methylating sites (Ext. Data Fig. 5), and in a sensitivity analysis in which we omitted the initial trimming of ages up to the age of sexual maturity (Extended Data Fig. 6). Notably, the three largest outliers in blood samples were livestock species (pig, sheep, and cattle).

Overall, our analysis of DNA methylation data in mammals reveals scaling between maximum lifespan and DNA methylation rate over approximately two orders of magnitude and in two distinct tissue types. For blood, for example, this relationship means that the methylation rate of humans is about half as fast as that of chimpanzees, given that our lifespan is about twice as long. An interesting application of such scaling relationships is that it allows estimations of maximum lifespan for newly discovered species through longitudinal sampling, where only the time interval between samples is needed instead of any knowledge of chronological ages.

Many physiological characteristics exhibit scaling with lifespan because they are indirectly associated through body mass^4,17^. However, the relationship we observed is largely independent of body mass, with no clear trend seen when regressing against mass instead of lifespan (Extended Data Fig. 7). For example, the naked mole rat, which is an outlier in body mass relationships, scaled appropriately in our data^18^.

The fact that specific and quantitative relationships exist between methylation rate and maximum lifespan suggests that there is an evolutionary constraint acting across diverse mammalian lineages. This means that when an organism’s lifespan evolves, its methylation rates also change. A scaling law emerges when these changes occur in a predictable way.

Methylation changes over a lifespan can be most simply described by the occurrence of epimutations in stem cells and their inheritance through cell divisions. As such, methylation rate in a mammal, *M*_*L*_, of lifespan *L* can be thought of as the product of two underlying quantities: *R*, the rate of stem cell division, and *p*, the probability that a cell division results in a change in methylation state^19,20^.

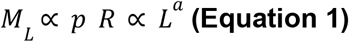

where *a* denotes the scaling law. Under this model, the quantity *pR* must scale with lifespan. As for which of these factors may be responsible for the scaling we observe, we discuss two non-mutually exclusive scenarios below.

Firstly, the probability of methylation changes with each stem cell division, *p*, may scale with maximum lifespan. In this scenario, aberrant methylation levels themselves are an evolutionary constraint on maximum lifespan. In other words, epimutation burden is deleterious and so mechanisms to reduce it are selected for in longer-lived organisms (i.e. *p* decreases as *L* increases in **Eq. 1**). This scenario would support an instructive role of DNA methylation in the associations observed between epigenetic changes and physiological outcomes in ageing^20–23^.

Secondly, it is possible that stem cell replication rates scale with maximum lifespan (i.e. *R* decreases as *L* increases in **Eq. 1**). In the haematopoietic system, it has previously been suggested that the rate of stem cell divisions in mammals decreases with lifespan such that the total number of divisions per stem cell is approximately constant, regardless of lifespan^24^. Additionally, it has recently been observed by Cagan et al. that somatic mutation rates also exhibit negative scaling with maximum lifespan^25^. Given that cell division plays a role in both processes, one possibility is that the scaling of both methylation and mutation rates is driven by stem cell replication rates.

While our study unambiguously shows the existence of scaling between methylation rates and maximum lifespan in mammals, the precise value of the scaling is subject to some uncertainty. Firstly, the data is one source of error. Specifically, although the Mammalian Methylation Consortium dataset provided an unprecedented resource for the community, the number of observations per mammal was sometimes limited. Additionally, the sampling distribution of ages in some mammals was uneven, sparse, or covered only a small proportion of their maximum lifespan. Combined, these factors addedd uncertainty to the calculation of methylation rates. Similarly, the age and maximum lifespan estimates in some mammalian species are imprecise, adding further uncertainty to the calculations. Second, while it is well-established that methylation dynamics can be approximated by linear functions, they do not provide a comprehensive model. Some age-associated CpGs exhibit non-linear dynamics later in life as they approach a stable value and begin to plateau. This phenomenon would result in an underestimation of the true methylation rate in faster-methylating mammals and likely bias our results towards the null (Extended Data Fig. 8) - although we partially mitigated this phenomenon by excluding CpGs with methylation values concentrated near the boundaries. Given all of these limitations, there is some uncertainty around the exact value of the scaling laws in blood and skin, and it is unclear whether they are distinct or converge on a common value. However, regardless of the precise values, it is striking that such strong scaling relationships exist even with all of these potential sources of error. Future studies could evaluate whether these scaling relationships hold for other classes and tissues and whether non-linear models may shed more light on the precise values of the scaling laws.

## Methods

### Considerations in the calculation of methylation rates

In contrast to previous studies^10,11^, we restricted our analysis to CpGs which were age-related in each mammal being compared. We did this because even a conserved CpG site may behave markedly differently between species. For example, the ELOVL2 CpG (cg16867657) is very strongly age-associated in humans and other primates but shows no age-association in most other species (Extended Data Fig. 9). As such, using this CpG to compare methylation rates between a primate and non-primate species may not be appropriate.

Additionally, we compared species across the same timespans because various statistical issues may arise when comparing methylation rates of species across different age ranges. In our study, there were two main considerations. Firstly, use of an R^2^ threshold (or equivalent) to select age-related sites may bias results towards slower rates in longer-lived animals. This is because slowly methylating sites may be unable to be detected in shorter-lived animals due to smaller timespans and the fact that methylation data is often noisy. In other words, shorter-lived animals might not be sampled long enough for small trends to become statistically apparent. Secondly, methylation proportions are bounded at 0 and 1. This is important as it means that if any given site is age-related, then methylation levels may be more likely to have approached these boundaries and begun to plateau in longer-lived mammals. If this is the case, then simply fitting linear regression lines to this data would result in slower methylation rates for longer-lived animals even with the same underlying dynamics (Extended Data Fig. 2).

Our initial approach was to find age-related CpGs common to all species and compare the average slope. However, very few CpGs satisfied this criterion, resulting in unstable estimates. This method would also extend poorly to additional animals as the number of common CpGs would decrease with each addition. Further, we were not able to compare animals over the same timespan given the vastly different sampling ranges between the shortest and longest-lived animals.

Because of all the above reasons, we decided to compare each species in a pairwise manner. This maximised the number of common age-related sites we could use in each comparison. Additionally, if we first ordered our dataset by maximum observed age, we could compare neighbouring species across the same timespan by appropriately trimming the datasets in each comparison. This would ensure a fair comparison while maximising the amount of data retained in each comparison. Finally, as we can sequentially move across the dataset, at each point calculating how the rates of the next mammal compare to the one before it, we can (indirectly) compare the methylation rates of the shortest-sampled animals with the longest, while at all points comparing rates over the same timespans.

### Primary analysis

We aimed to compare methylation rates - defined as the slope from linear regressions of methylation ∼ age - in conserved age-related sites across mammals. An overview of our workflow is depicted in Fig. 1a. We initially curated our data for each tissue by conducting principal component analysis on all species combined. We then projected each tissue onto the PC1 and PC2 components to detect and remove outlier samples using density-based spatial clustering (DBSCAN) for each species separately (Step 1 in Fig. 1a, Extended Data Fig. 1, see code for further details). Additionally, we removed samples below the age of sexual maturity known to exhibit non-linear dynamics^7^ (Step 2 in Fig. 1a). The age of sexual maturity, as reported in the AnAge database^26^, was then subtracted from all ages so that 0 represented the age of sexual maturity.

To develop a statistically robust framework we therefore ordered the datasets by maximum observed age and compared each mammal with its neighbours in a sequential pairwise manner (Step 3 in Fig. 1a and Methods). We started from the mammal with the shortest observed age, which we refer to as the baseline species. For each comparison we restricted the datasets of both mammals to be as close as possible to each other (Step 4 in Fig. 1a and Methods). Specifically, we calculated the maximum sample age of the shorter-observed species and then found the sample with closest age from the comparsion species (either above or below the maximum sample age of the shorter-observed species). If the differences in samples was greater than 2% of the lifespan of the shortest mammal or 1 year (whichever was smallest), we used the next-oldest sample in the shorter-observed species and repeated the process. Once two ages were found that satisfied these criteria, we then restricted the observations in each of the compared species accordingly. Mammals which had fewer than 15 samples after this restriction (or 20 initially) were excluded, as were species for which the maximum sampled age was below 25% of the reported maximum lifespan.

For each pairwise comparison, we selected the common set of age-related CpGs passing an R^2^ threshold and sharing the same directionality. A CpG was considered age-associated if the R^2^ value from a simple linear regression passed a certain threshold (see Additional Analyses). CpGs with a mean methylation proportion above 0.9 or below 0.1 (calculated after any data trimming) were removed as these sites tend to display non-linear dynamics due to being near the maximum/minimum methylation values.

For each CpG satisfying these criteria in both mammals in each pairwise comparison, we calculated the methylation rate for each species (Step 5 in Fig. 1a). We then calculated the methylation rate ratio of the longer-sampled species compared to the shorter-sampled species. That is, for each CpG, methylation rate ratio = rate of longer-sampled species/rate of shorter-sampled species. For example, a ratio of 0.5 would mean that the methylation rate of the longer-sampled species is twice as fast as the methylation rate of the shorter-sampled species in a particular CpG. We calculated these ratios across all CpGs included for each comparison and extracted the median ratio. Use of the median was chosen over the mean as the mean was severely affected by large outliers (resulting from the division of very small numbers in some ratio calculations).

Then, we computed the cumulative product of median ratios to compare all species together (Step 6 of Fig. 1a, and Methods). This cumulative product can be thought of simply as the methylation rate of each species (indirectly) compared to the baseline species. For example, the first mammal in the list (e.g. rat) is given a rate ratio of 1. Suppose the next mammal in the list (mouse) is compared to the rat, yielding a rate ratio 0.94 (i.e. its median methylation rate is 0.94 as fast as that of a mouse, in common conserved age-related CpGs). Suppose the next mammal in the list (sheep) is then compared to the mouse, yielding a rate ratio of

0.53. We can indirectly compare the rates of the miniature pig to those of the mouse by the cumulative rate ratio of 0.94*0.53=0.49, and so on. In this way, we can compare the methylation rates of the shortest-sampled animals with the longest, while at all steps comparing rates over the same timespans.

We include a more detailed mathematical description of this process below.

### Mathematical description of scaling and power laws

Mathematically, scaling between two variables *x* and *y* can generally be described as a power law relationship of the form

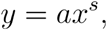

where *s* is the scaling law and *a* is a constant. Alternatively, taking the logarithm on both sides allows for linear inference of both *s* and *a*,

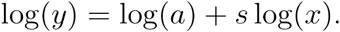

In our case, we are interested in inferring the scaling law relating the lifespan, *l*_*m*_, of a mammal, *m*, with the slope, *S*_*c,m*_, of the methylation values in a CpG site *c*. This translates to the following scaling relation

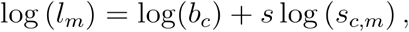

where *b*_*c*_ denotes the baseline slope in site *c*, or slope predicted for a mammal with a lifespan of 1 year.

Our pairwise comparison algorithm exploits the following relation, for any 2 mammals *m*_0_ and *m*_1_,

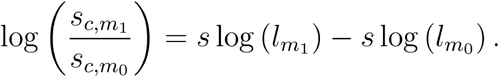

In other words, the ratio between the slopes in two mammals replaces the intercept in the linear relation with one that is relative to the lifespan of the baseline species *m*_0_.

Finally, given an arbitrarily ordered set of species *m*_0_, *m*_1_, … *m*_i_, the cumulative product of slopes results in a estimators of the desired scaling law *s* :

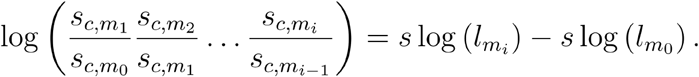

### Stability of scaling law

We calculated the impact of varying the minimum R^2^ threshold to define an age-related CpG (see Extended Data Fig. 3). A grid of R^2^ values between 0 and 0.2 were explored for each tissue (see Extended Data Fig. 3a, 3c). We then assessed the stability of our results using the kernel density estimate of all reported values and selected the optimal scaling as the point of maximum density. (see Extended Data Fig. 3d, 3d). R^2^ values that resulted in less than 10 CpGs in any one comparison were not considered, even if below the 0.2 threshold.

### Biases and statistical robustness

We conducted various analyses to explore the robustness of our results.

Firstly, we conducted a random null simulation, showing that not accounting for differences in sample age ranges results in an artificial negative association with maximum lifespan (Extended Data Fig. 2b). Specifically, we created synthetic data representing a scenario that was as similar as possible to the real data, except with no scaling of methylation rates. We used the observed species and associated lifespans in our datasets, but with synthetic methylation data. For each species and site we uniformly sampled ages within its lifespan and randomly drew slopes and initial methylation values from normal distributions to create linear data with random noise and constrained its values to within 0 and 1. In this analysis, we simply calculated the methylation rates of each mammal across their entire sampled age ranges.

Secondly, we again used synthetic data, but instead conducted our primary analysis method (Fig. 1a) on it. In this case, we used the observed species and used exact sampled ages in our datasets but randomly drew slopes and intercepts from all those observed across all mammals in this study to create synthetic linear data.

## Data availability

The majority of the methylation dataset used was created by the Mammalian Methylation Consortium^27^ and is publicly available on the Gene Expression Omnibus at accession number GSE223748. The chimpanzee (*Pan troglodytes*) dataset is available at GSE136296^28^.

Data on maximum lifespan and mass were taken from the AnAge database^26^.

## Code availability

All code used for the analysis (conducted in Python 3.11.4) is available at https://github.com/elc08/meth_scaling_law.

## Acknowledgements

We thank Thomas Little for critical reading of the manuscript, and Melike Dönertaş for bringing our attention to additional statistical considerations. We thank all members of the Chandra lab for their input. S.J.C.C is supported by the Wellcome Trust Hosts, Pathogens & Global Health Programme (grant number: grant.226831/Z/22/Z). E.L.C. is a cross-disciplinary postdoctoral fellow supported by funding from the University of Edinburgh and the Medical Research Council (MC_UU_00009/2). T.C. is supported through a Chancellor’s Fellow at the University of Edinburgh and the MRC Human Genetics Unit.

## Extended data

**Extended Data Fig. 1:**
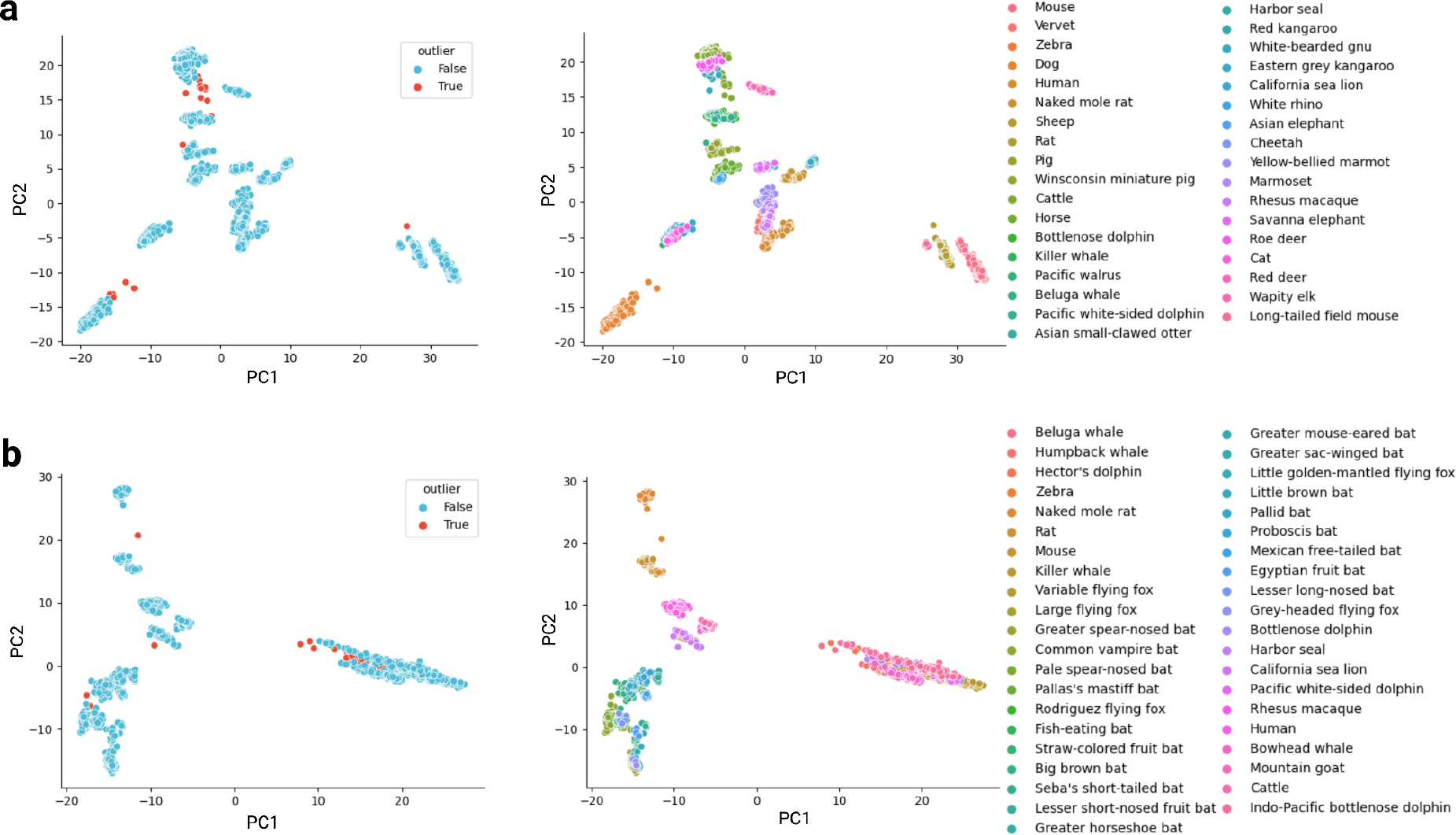
PCA-based outlier removal. **a**, Left: Result of outlier detection using the density-based spatial clustering of applications with noise (DBSCAN) algorithm on PC1 vs PC2 plots resulting from principal component analysis (PCA) of all blood samples combined. Right: Same as the lefthand plot, but coloured by species instead of outlier status. **b)** Same analysis as in a), except on skin samples.

**Extended Data Fig. 2:**
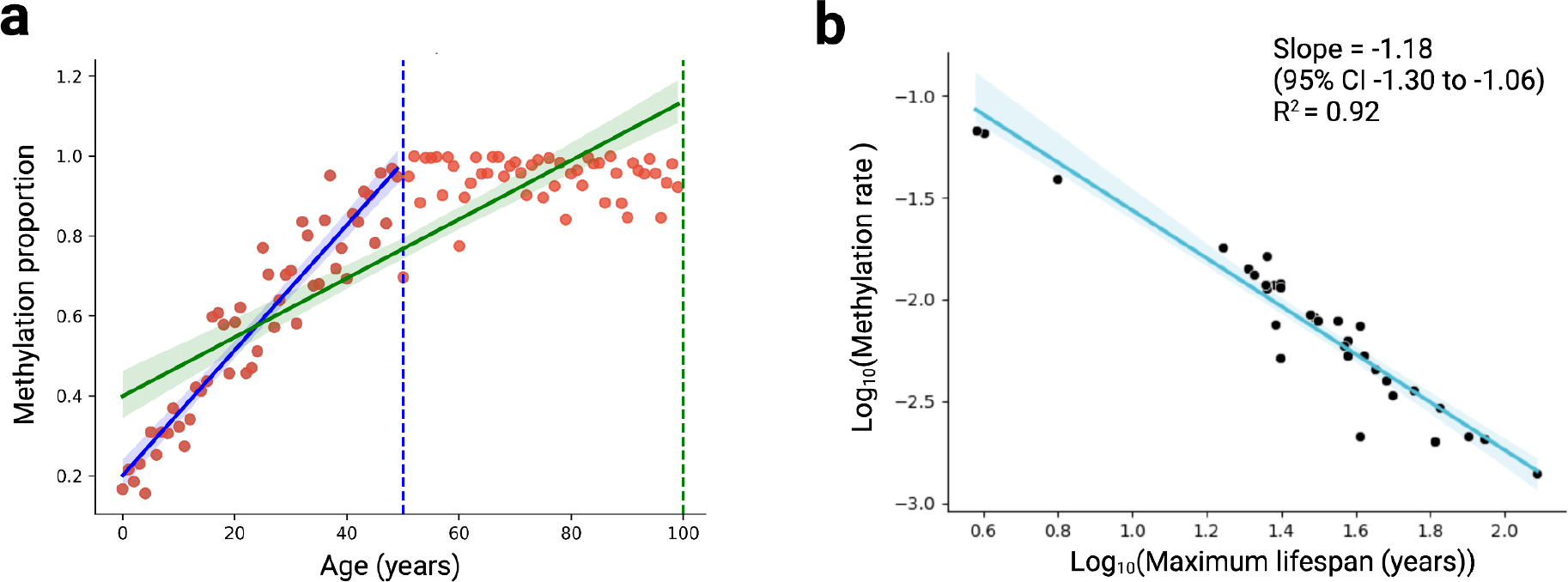
Potential statistical issues arising when comparing methylation rates across different timespans. **A**, Differences arising in fitting a linear slope to identically generated synthetical data in a shorter-lived mammal that is sampled to a certain age (50 years, dotted blue line) and a longer-lived mammal sampled for a longer time (100 years, dotted green line). Linear fits and 95% confidence intervals are shown in their respective colours. The estimated slope for the longer-lived species is under-estimated as the result of constraints in methylation range. **b**, Results from a random simulation, demonstrating that not accounting for differences in sample age ranges results in an artificial negative association between methylation rate and maximum lifespan. Each point corresponds to the median slope inferred in synthetically generated data for all mammals included in our analysis of the blood tissue (see Methods). Regression line from a linear regression of the form y∼x is shown in blue with its associated shaded region representing the 95% confidence interval.

**Extended Data Fig. 3:**
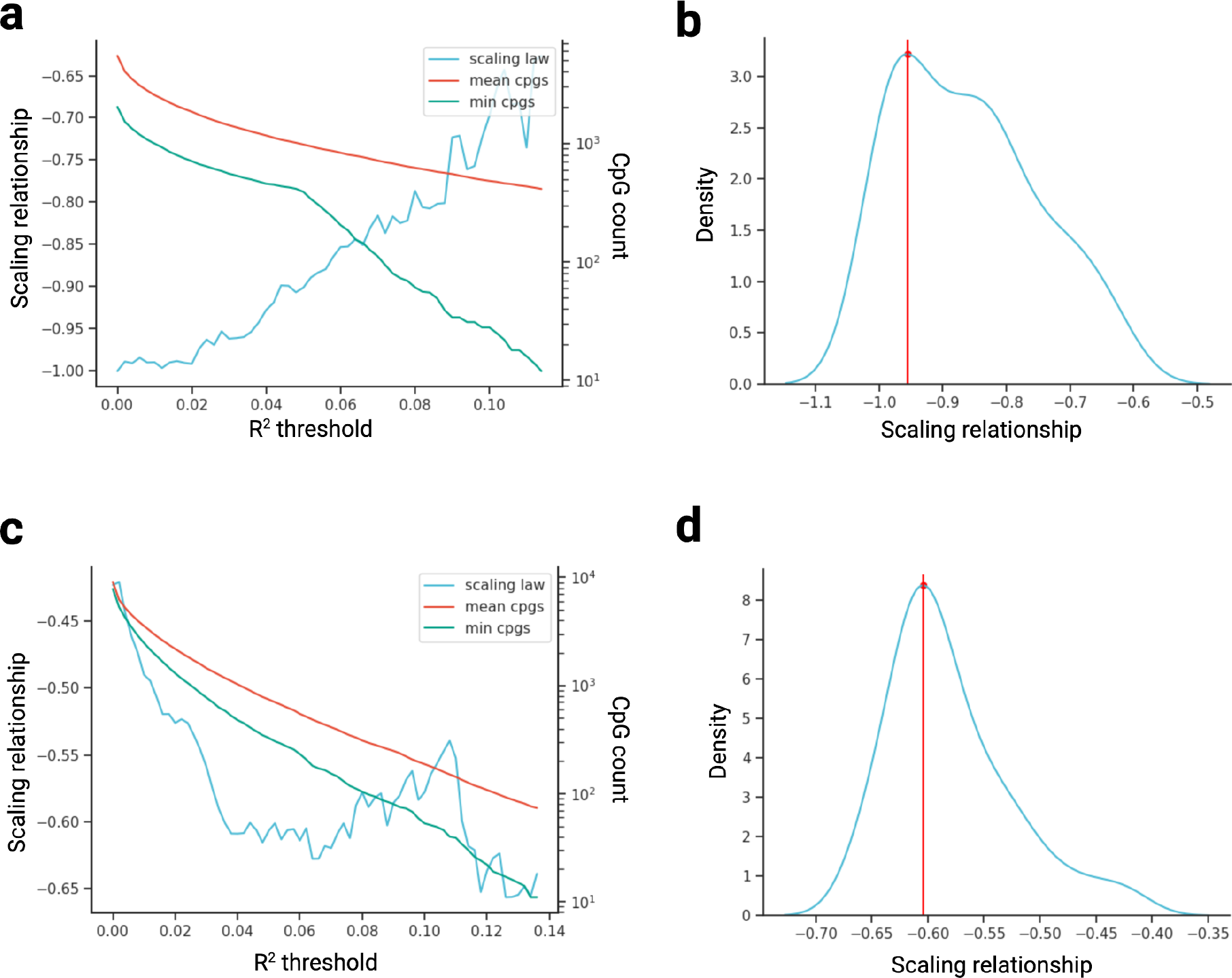
Scaling relationships across varying R^2^ thresholds used to define an age-related site. **a**, Scaling relationship (i.e. slope of the log-log plot of methylation rate ratio versus maximum lifespan) across varying R^2^ thresholds used to define an age-related site, in blood samples. Also shown is the mean number of CpGs used across all pairwise comparisons, and the minimum number of CpGs used in any single comparison. **b**, Kernel density estimate (KDE) plot of the scaling relationships shown in a). Red line shows the maximum density value. **c, d**, Same as in a) and b) except in skin samples.

**Extended Data Fig. 4:**
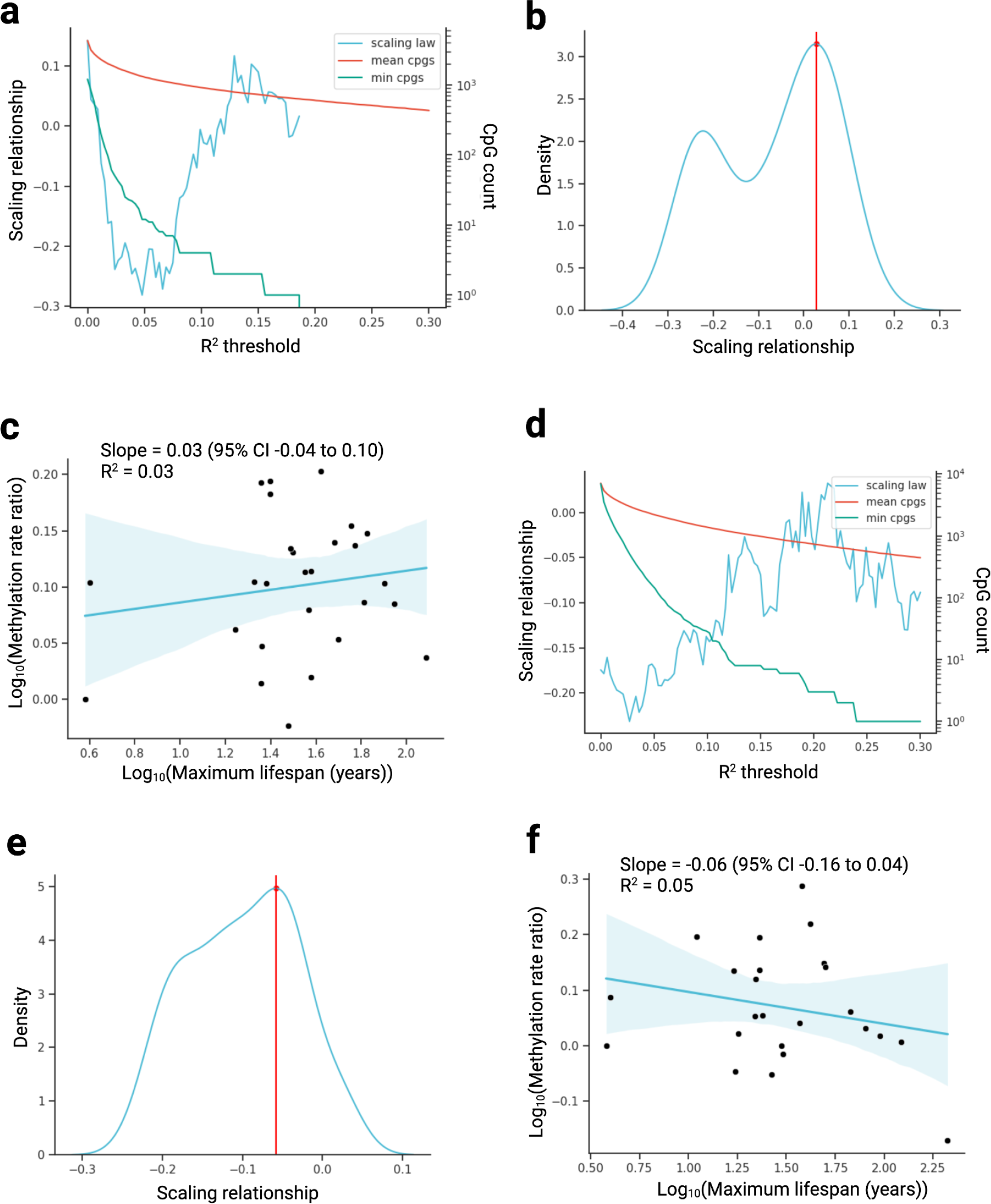
Scaling relationships using simulated null data. **a**, Scaling relationship (i.e. slope of the log-log plot of methylation rate ratio versus maximum lifespan) across varying R^2^ thresholds used to define an age-related site, in a random null simulation using blood samples. Also shown is the mean number of CpGs used across all comparisons, and the minimum number of CpGs used in any single comparison. **b**, Kernel density estimate (KDE) plot of the scaling relationships shown in a). Red line shows the maximum density value. **c**, Log methylation rate (ratio compared to baseline species) versus log maximum lifespan, with R^2^ threshold used to define an age-related site taken from the maximum density value in b) (red line). The y-coordinate for each point shows the log_10_ cumulative product of the median slope ratio (see Methods). Regression line from simple linear regressions of the form y∼x. Shaded region represents the 95% confidence interval. **d-f**, Equivalent analysis as in a-c, but in skin samples.

**Extended Data Fig. 5:**
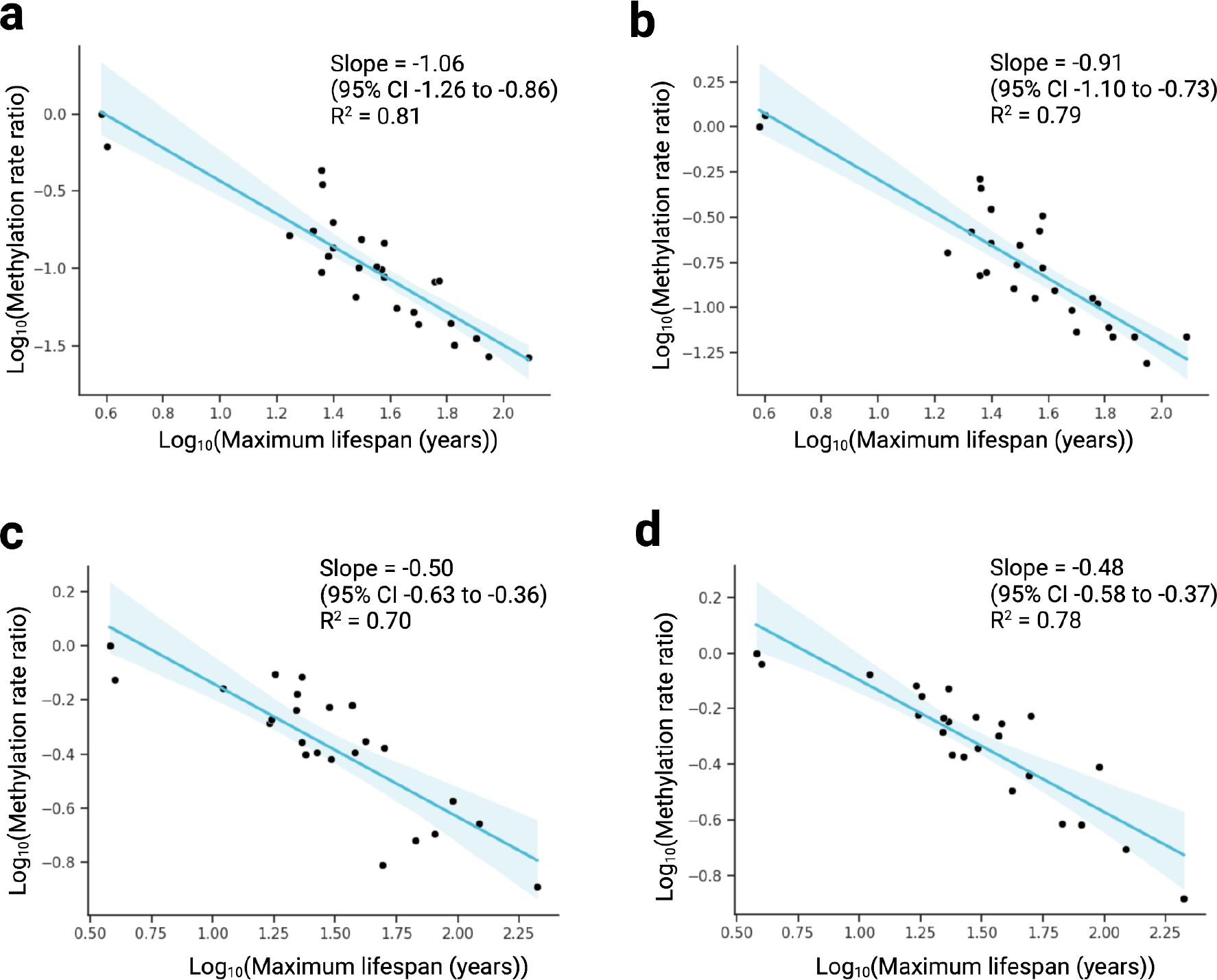
Scaling relationships stratified into hyper- and hypomethylating sites. **a**, Log methylation rate ratio, compared to baseline species, versus log maximum lifespan in blood samples in hypermethylating (i.e. increasing) sites only. The y-coordinate for each point shows the log_10_ cumulative product of the median slope ratio (see Methods). Regression line from simple linear regression of the form y∼x. Shaded region represents the 95% confidence interval. **b**, Equivalent analysis as in a, but in hypomethylating sites only. **c**,**d** Equivalent analysis as in a and b, but in skin samples.

**Extended Data Fig. 6:**
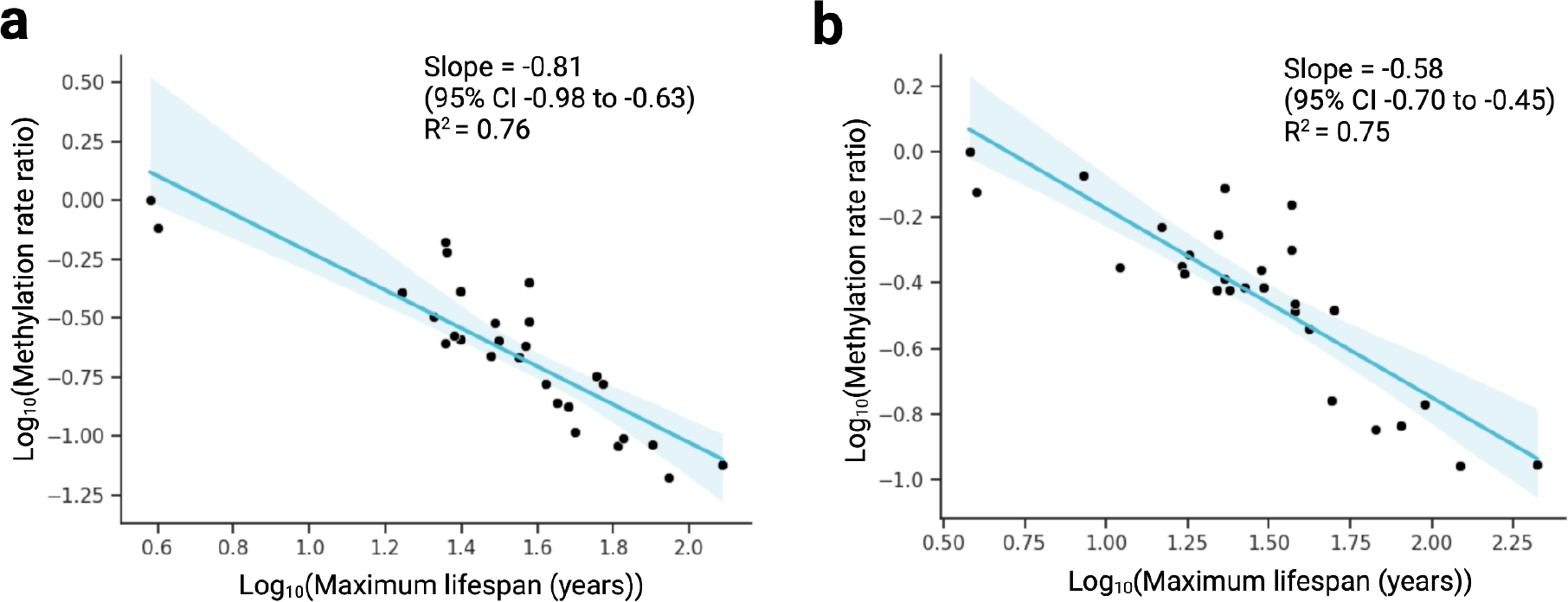
Scaling relationships without initial trimming of ages below sexual maturity. **a**, Log methylation rate ratio, compared to baseline species, versus log maximum lifespan in blood samples. Methylation rates are calculated for each species without initial trimming of ages below sexual maturity. The y-coordinate for each point shows the log_10_ cumulative product of the median slope ratio (see Methods). Regression line from simple linear regression of the form y∼x. Shaded region represents the 95% confidence interval. **b**, Equivalent analysis as in a), but in skin samples.

**Extended Data Fig. 7:**
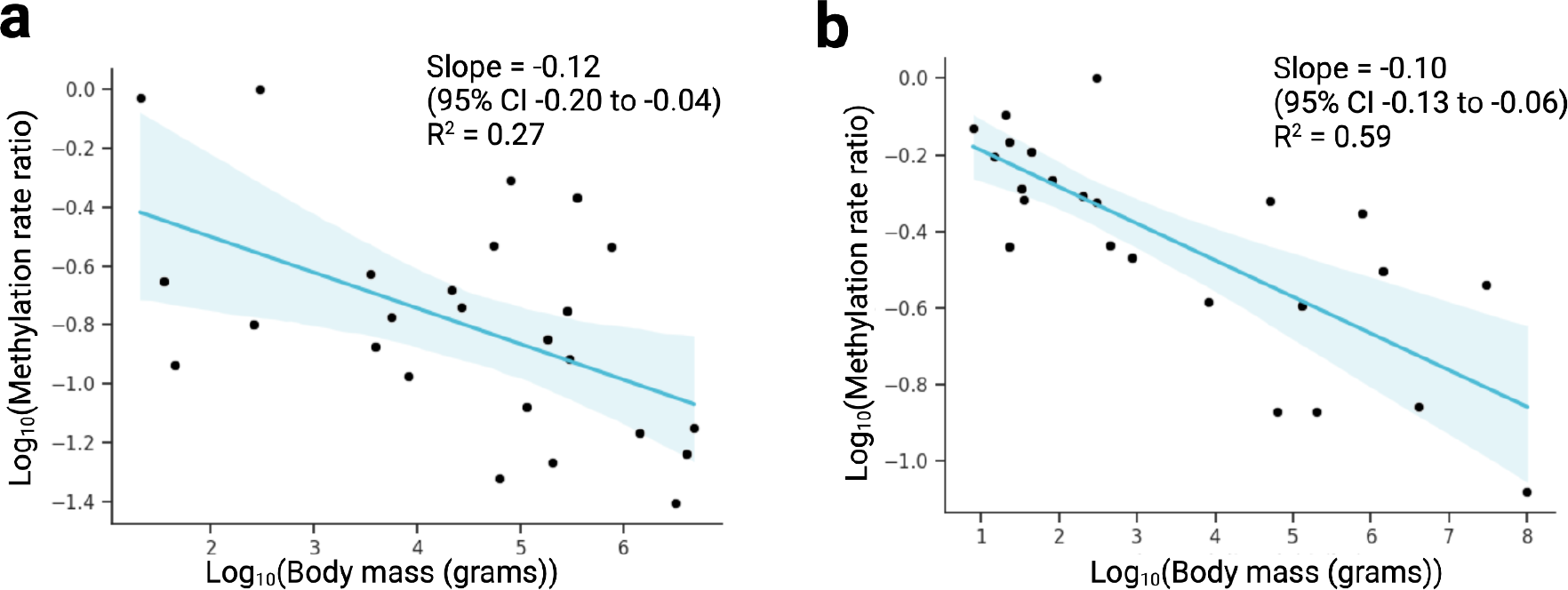
Scaling relationship between methylation rate and body mass. **a**, Log methylation rate (ratio compared to baseline species) versus log body mass in blood samples. The y-coordinate for each point shows the log_10_ cumulative product of the median slope ratio (see Methods). Regression line from simple linear regression of the form y∼x. Shaded region represents the 95% confidence interval. **b**, Equivalent analysis as in a, but in skin samples.

**Extended Data Fig. 8:**
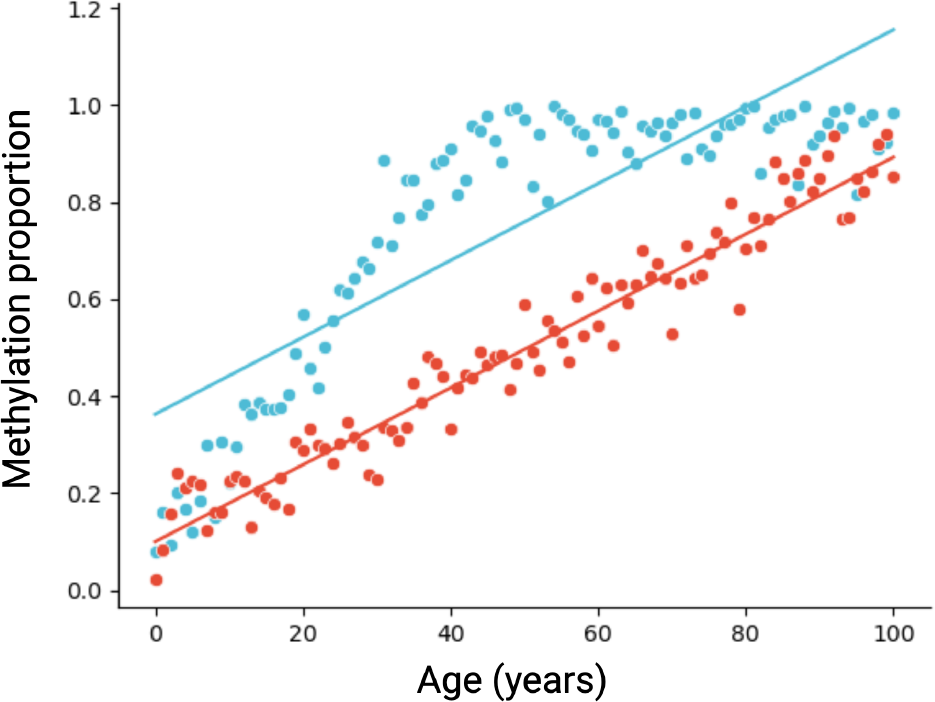
Potential remaining bias, leading to an underestimation of methylation rates of faster-methylating species. Synthetic data showing a faster methylating site (blue) that reaches the maximum methylation value (1) and begins to plateau, compared to a slower methylating site (red) that does not plateau. In each case, the resulting inferred methylation rates (represented by the linear regression lines) are identical.

**Extended Data Fig. 9:**
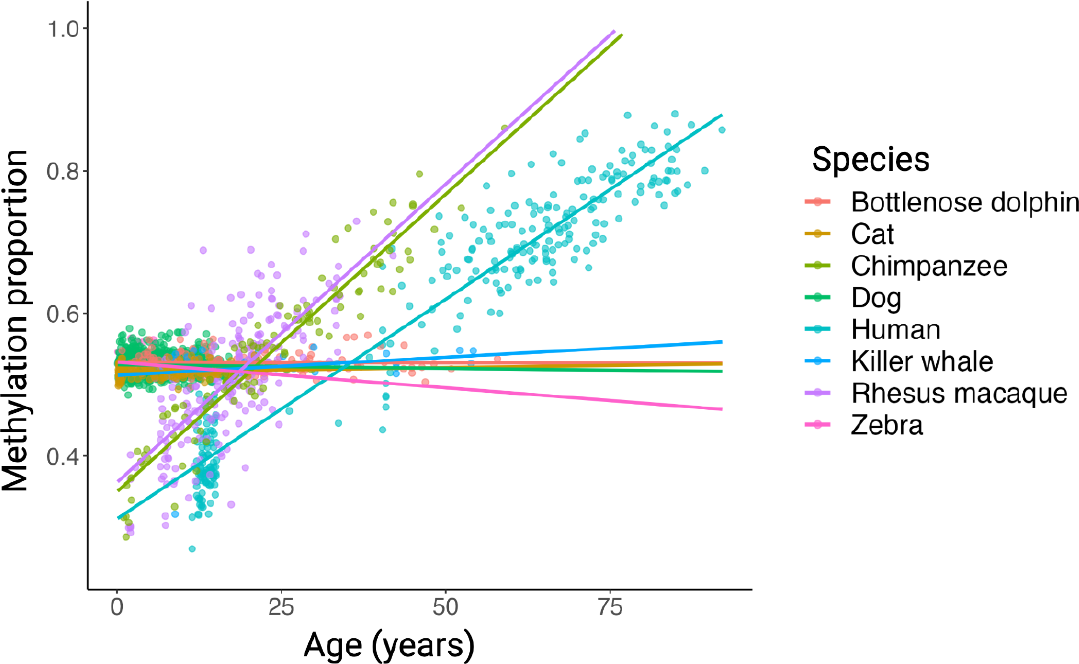
ELOVL2 CpG. Example of a CpG (ELOVL2) that is strongly age-related in some species but relatively age-invariant in others. Regression lines from simple linear regressions of the form methylation ∼ age.

### Description of extended data tables

**Extended Data Table 1:** Details of mammals used in the primary analysis of blood samples. Columns describe common name, lifespan (years), and average adult weight (grams).

**Extended Data Table 2:** Details of mammals used in the primary analysis of skin samples. Columns describe common name, lifespan (years), and average adult weight (grams).

**Extended Data Table 3:** Results of the primary analysis in blood samples. Each row describes a pairwise comparison. Columns describe: reference mammal, comparison mammal, maximum age (with zero representing sexual maturity) of the reference mammal after trimming of sample age ranges, maximum age (with zero representing sexual maturity) of the comparison mammal after trimming of sample age ranges, sample size of the reference mammal after trimming of sample age range, sample size of the comparison mammal after trimming of sample age range, median slope ratio in the pairwise comparison, and cumulative median slope ratio (i.e. ratio compared to the baseline species).

**Extended Data Table 4:** Results of the primary analysis in skin samples. Each row describes a pairwise comparison. Columns describe: reference mammal, comparison mammal, maximum age (with zero representing sexual maturity) of the reference mammal after trimming of sample age ranges, maximum age (with zero representing sexual maturity) of the comparison mammal after trimming of sample age ranges, sample size of the reference mammal after trimming of sample age range, sample size of the comparison mammal after trimming of sample age range, median slope ratio in the pairwise comparison, and cumulative median slope ratio (i.e. ratio compared to the baseline species).

